# Mucosal vaccination for SARS-CoV-2 elicits superior systemic T central memory function and cross-neutralizing antibodies against variants of concern

**DOI:** 10.1101/2022.09.09.507250

**Authors:** Aled O’Neill, Monica Palanichamy Kala, Tan Chee Wah, Wilfried A.A. Saron, Chinmay Kumar Mantri, Abhay P.S. Rathore, Lin-Fa Wang, Ashley L. St. John

## Abstract

COVID-19 vaccines used in humans are highly effective in limiting disease and death caused by the SARS-CoV-2 virus, yet improved vaccines that provide greater protection at mucosal surfaces, which could reduce break-through infections and subsequent transmission, are still needed. Here we show that intranasal (I.N.) vaccination with the receptor binding domain of Spike antigen of SARS-CoV-2 (S-RBD) in combination with the mucosal adjuvant mastoparan-7 improved systemic T cell responses compared to an equivalent dose of antigen delivered by the sub-cutaneous (S.C.) route, adjuvanted by either M7 or the gold-standard adjuvant, alum. T cell phenotypes induced by I.N. vaccine administration included enhanced polyfunctionality (combined IFN-γ and TNF expression) and greater numbers of T central memory (T_CM_) cells. These phenotypes were T cell-intrinsic and could be recalled in the lungs and/or brachial LNs upon antigen challenge after adoptive T cell transfer to naïve recipients. Furthermore, mucosal vaccination induced antibody responses that were similarly effective in neutralizing the binding of the parental strain of S-RBD to its ACE2 receptor, but showed greater cross-neutralizing capacity against multiple variants of concern (VOC), compared to S.C. vaccination. These results highlight the role of nasal vaccine administration in imprinting an immune profile associated with long-term T cell retention and diversified neutralizing antibody responses, which could be applied to improve vaccines for COVID-19 and other infectious diseases.

## Introduction

SARS-CoV-2 emerged in 2019 as a novel Coronavirus infecting humans and in 2020 it began a major ongoing global pandemic. The disease it induces in humans, COVID-19, is characterized by fever, cough, fatigue and dyspnea, with severe cases leading to pneumonia and death^1^. Vascular complications and coagulation disorders also occur^2^. The elderly and those with pre-existing conditions, such as diabetes and hypertension, are most at risk for developing life-threatening complications^1,3^. The widespread worldwide distribution of active COVID-19 infection clusters and the severity of disease outcomes in patients in multiple age groups has necessitated unprecedented advances in vaccine technologies and distribution. Although a multitude of vaccines are now available that show protection in terms of significantly reducing the incidence of infections, hospitalizations, deaths and reducing transmission^4-8^, breakthrough infections often occur^9^, suggesting that there are limitations to the duration of protective immune responses induced by the current vaccine regimens. Furthermore, new variants of concern (VOC) continue to circulate, even in populations with high levels of vaccine coverage^10^. This is thought to be at least partially attributable to immune pressure on the SARS-CoV-2 virus leading to diversification of antigenic properties through virus mutation^10^.

Among the first vaccines approved against SARS-CoV-2 were mRNA-based vaccines, which initial analyses showed can be >90% effective a few weeks following the vaccine protocol completion^6,11^. Most strategies have used the Spike (S) protein as antigen, which is found on the virus surface. Often, the region of the S-protein containing its receptor binding domain (RBD) that allows its entry into host cells via binding to the angiotensin-converting enzyme 2 (ACE2) receptor is used. Importantly, ACE2 is expressed by the type I and II alveolar cells of the lung that are key targets of lower respiratory tract infection by SARS-CoV-2^12-14^, so that neutralizing antibodies against this protein are effective in preventing cellular entry and infection by the virus. For human vaccinees who were given mRNA vaccines, there are strong correlations between the titer of vaccine-induced antibody responses and protection from symptomatic disease^15^. Notwithstanding their efficacy, risk of breakthrough infection appeared to increase in the months following completion of the two-dose mRNA vaccine^9^, likely owing to the natural time-related decay in specific antibodies. Complicating this phenomenon of waning protection over time has been the emergence of VOC, for which vaccine-induced antibodies show decreased neutralization^10^. Despite the loss of antigen-specificity and neutralization capacity of vaccine-induced antibodies to VOC such as Omicron^16^, vaccinees remain highly protected against severe disease and death^17^, which could possibly point to a protective role for T cells in vaccine-induced protection. Indeed, T cells are highly cross-reactive to VOC and even to SARS-CoV-1 and seasonal coronaviruses^18-20^. In primates with SARS-CoV-2 infection, CD8 T cell activation correlated with viral control in the absence of neutralizing antibodies^21^. In humans, rapid induction of SARS-CoV-2-specific T cell responses were also associated with mild disease^22,23^. Boosting of mRNA vaccines has been shown to lead to a surge in vaccine protection that correlated with the boost in S-specific antibody titers^24^. These observations highlight outcomes of COVID-19 vaccines that could be further improved as our understanding of functional correlates of COVID-19 protection grows.

mRNA vaccines also have limitations for world-wide use given that they are difficult to distribute and require strict cold chain adherence and storage near -80°C^25^. Several alternative vaccine approaches are also being developed for SARS-CoV-2, including subunit vaccines^26^, which involve use of more-stable protein antigens and have an advantage for stability at multiple temperatures. Although subunit vaccines have been used effectively in the context of many viral vaccines, including those approved for hepatitis B and influenza viruses^27,28^, usually, the protein components of subunit vaccines are not sufficient, alone, to establish immune memory^29^. For this reason, adjuvants, or substances that promote immune activation, are often added to the subunit vaccine formulation to induce long-term memory responses. Alum and AS04 (aluminium salt combined with the TLR4 agonist 2-O-desacyl-4’-monophosphoryl lipid A) are two human-approved adjuvants, but these are not used at mucosal surfaces^30,31^. Currently, there are no mucosal adjuvants approved for use in humans, but there are adjuvants that have been used in mucosal vaccine formulations in experimental settings, including the most widely studied experimental mucosal adjuvant, Cholera toxin^32^, and mast cell activating compounds, such as mastoparan^32-34^. Cholera toxin causes toxicity so it cannot be used in humans^32^. Mastoparan is a short 14aa peptide that is of insufficient length to trigger immune responses itself. Its analogue, mastoparan-7 (M7) has greater cell-activating activity and appears to work in vivo primarily through inducing mast cell degranulation responses through the MrgX2 receptor^35^. M7 is effective in enhancing the titer of antigen-specific antibodies in animal models when delivered in combination with vaccine antigens, both through S.C. injection as well as application to the nasal mucosae^36^. In a haptenated cocaine vaccination strategy, M7 augmented antibody responses that prevented the psychoactive effects of cocaine. This was likely through its enhanced titers of IgA and improved antigen-specific IgG avidity compared to similar subcutaneous vaccinations with Alum used as adjuvant^34^. However, it is unknown if M7 works differently in the skin compared to mucosal sites, induces site-specific responses, or influences T cell phenotypes and functions that are particularly important for combatting certain types of viral pathogens. Prior studies also established the adjuvant activity of M7 during homologous challenges, but whether mucosal vaccines generally or M7-adjuvanted vaccines specifically induce responses that are more broadly protective against diverse viral isolates compared to conventional approaches is also unknown. For vaccines against SARS-CoV-2, mucosal vaccines have shown promise in experimental studies, with multiple platforms, including unadjuvanted S protein and viral vector-based systems evoking protective immune responses^37-42^.

Given that SARS-CoV-2 infection is initiated at the mucosal surface of the nasal passages and lung airways, we planned this study with the aim of testing whether delivery of an adjuvanted subunit vaccine intra-nasally (I.N.) has advantages for the induction of SARS-CoV-2-specifc and protective immune responses. We found that mucosal administration of adjuvanted SARS-CoV-2 subunit vaccines is a promising strategy to improve systemic immune responses, through preferential induction of central memory T (T_CM_) cells that are polyfunctional. These T_CM_ responses are T cell intrinsic and are maintained following transfer to new hosts to promote improved memory recall upon lung antigen challenge in both the draining brachial lymph nodes (LNs) and lungs. Furthermore, the improved polyfunctional response following intra-nasal (I.N.) vaccination is extended to antibodies, which show improved breadth of neutralizing responses against multiple variants compared to vaccination subcutaneously (S.C.) with the same adjuvant.

## Results

### Superior systemic T cell responses with mucosal adjuvant

We began by comparing the ability of various vaccine formulations utilizing recombinant S protein to activate T cells. For this, mice were immunized S.C. with the receptor binding domain (RBD) of S, combined with either adjuvant, Alum or M7 (**Fig. 1A**), or I.N. with an equivalent amount of S-RBD and M7 (**Fig. 1B**). Although the RBD is small, our assessments suggested that there are multiple CD4 and CD8 T cell epitopes predicted for mice in this region of the S protein (**Table S1-2**), which is consistent with the observations in humans that the RBD contains confirmed T cell epitopes^43^. The T cell responses 5 days post-immunization were measured by flow cytometry in the draining lymphoid organs for the respective tissues for S.C. or I.N. immunizations, respectively, either the popliteal LN or the nasal-associated lymphoid tissue (NALT), the latter of which is the rodent structure analogous to Waldeyer’s ring in humans^44^. Systemic T cell responses were also assessed in the spleen following either route of immunization. The various populations of T cells found in the lymphoid organs were visualized using the UMAP algorithm (**Fig. 1C**) and were identified using the gating strategy shown in **Fig. 1D**. We first compared the numbers of total and activated T cells in these secondary lymphoid organs after subcutaneous or nasal injection with M7+S-RBD to saline-treated or S-RBD antigen alone-treated control groups. Subcutaneous immunizations with M7 induced increased retention of total CD3^+^ T cells as well as CD8^+^ and CD4^+^ T cell subsets in the draining popliteal LN (PLN), and conventionally innate T cells, γδ and NKT cells, followed a similar trend (**Fig. 1E, S1A-C**). Vaccination with alum+S-RBD also induced strong T cell activation responses in the local draining lymph node, similar to M7+S-RBD delivered S.C. (**Fig. 1E**). In contrast, the total T cell and CD8^+^ T cell numbers were not significantly affected in the NALT following mucosal challenge with M7+S-RBD, while there was a small but significant increase in the total number of CD4^+^ T cells (**Fig. 1E, S1A-C**). There were also differences in activated T cells, with increased numbers of total and CD69^+^CD4^+^ T cells in the NALT following I.N. vaccination, which was not observed in the PLN after S.C. vaccination (**Fig. 1E, S1D**).

**Figure 1:**
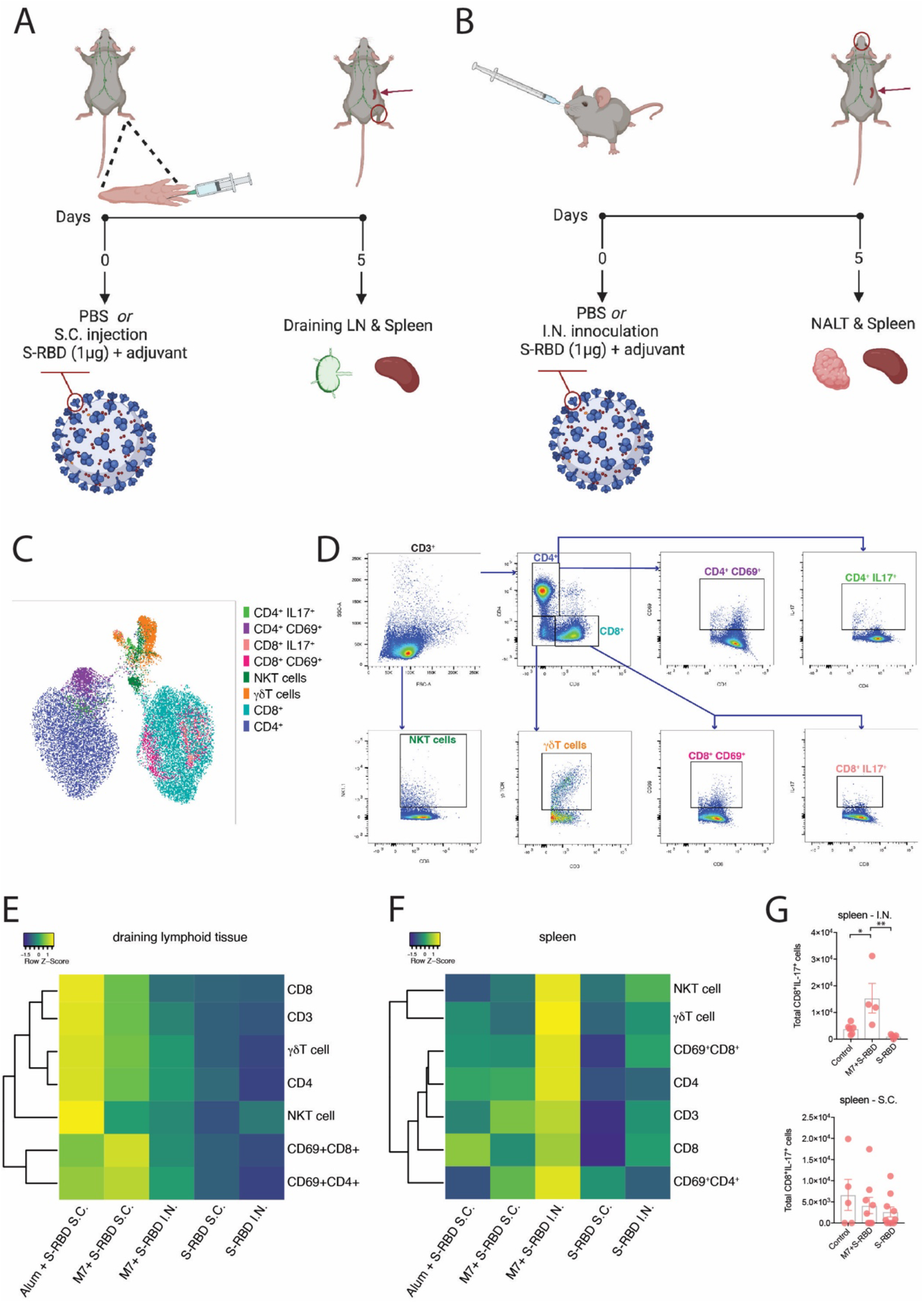
Superior systemic T cell activation following mucosal vaccination against SARS-CoV-2. (**A-B**) Diagrams of (**A**) sub-cutaneous (S.C.) and (**B**) intra-nasal (I.N.) vaccination strategies. (**C**) UMAP representation of the populations of T cells assessed in lymphoid organs 5 days following vaccination. (**D**) Flow cytometry strategy to identify T cells and their phenotypes and activation status corresponding to the populations depicted in panel C. (**E-F**) Heat map representations of the frequency of various T cell subsets in the (**E**) Draining lymphoid tissue (either NALT or popliteal LN) or (**F**) Spleen, day 5 following vaccination. The data in E-F are alternatively presented as graphs with error bars in **Figure S1**. (**G**) Numbers of IL-17^+^ CD8 T cells in the spleen following I.N. or S.C. vaccination with M7+S-RBD compared to controls or antigen (S-RBD) alone. *p<0.05 and **p<0.01 by 1-way ANOVA.

In contrast to the draining lymphoid organs, we noted that systemic activation of T cells was much higher in the spleen following mucosal vaccination. There were increased numbers of total T cells, as well as total, γδ, NKT and activated CD4 & CD8 cells after I.N. vaccination with M7+S-RBD compared to unvaccinated controls, which did not occur in the S.C. vaccinated groups (**Fig. 1F, S1A, S1F-I**). These data illustrate that mucosal immunization results in improved systemic immune activation, compared to peripheral sub-cutaneous immunization, even comparing the same adjuvant and antigen. In contrast, immune activation was more restricted to the draining lymphoid tissue following subcutaneous injection (**Fig. 1E-F**). These results highlighted a skewing of T cell activation in the draining lymphoid tissue towards increased CD4 activation in the NALT and increased CD8 activation in the spleen but, overall, support that immune activation is significantly induced in the draining lymphoid tissues after either S.C. or I.N. vaccination. I.N. vaccination was associated with stronger T cell activation in the spleen compared to S.C. vaccination.

In addition to T cell activation, we also measured intracellular cytokine expression in T cells, including IFN-γ, TNF and IL-17, as these cytokines define polarized T cell responses^45^. While intracellular IFN-γ and TNF were not detected at this time point, we observed IL-17^+^ expressing CD8^+^ T cells were uniquely enhanced in the spleen of mucosally-immunized mice, but not in the NALT of the same animals or in either the spleen or draining LNs of mice given S.C. immunizations with the same adjuvant, M7 (**Fig. 1G**). This is interesting given the fact that Th17 responses have been associated with IFN-γ -independent immune activation, augmented B cell activity, IgA induction at mucosal sites, and protective immune responses during respiratory viral infections^46^.

Given the strong systemic immune response observed in the spleen immediately following mucosal vaccination with M7+S-RBD, we next aimed to extend beyond the acute T cell responses elicited by the vaccine by comparing the antigen-specific T cell memory responses. For this, T cells were harvested from spleens 5-weeks post-vaccination for animals given S.C. versus I.N. challenges with the same antigen and adjuvant combination (M7+S-RBD) and tested *ex vivo* for their activation following antigen stimulation, according the experimental design shown in **Fig. 2A**. Memory T cell (T_MEM_) populations were identified with the gating strategy provided in **Fig S2B**. Antigen stimulation induced expansion and activation of CD4 and CD8 T effector memory (T_EM_) and T central memory (T_CM_) cells in vaccinated groups over the baseline found in naïve animals, but there were not significant differences in these populations between the two routes of vaccine administration (**Fig. S2B-H**), except for the elevated numbers of activated CD8 T_EM_ cells, based on CD69 expression, observed in the S.C. group (**Fig. 2B**). This seemed consistent with the strong induction of CD8 T cells at day 5 following S.C. immunization (**Fig. 1E-F**). Intracellular staining for cytokines identified a heightened functional response by both CD4 and CD8 T_CM_ cells, where a higher proportion of T_CM_ cells were TNF^+^ following I.N. compared to S.C. immunization (**Fig. 2C-F**). Furthermore, I.N. immunization induced more CD4 T_CM_ cells that were polyfunctional based on the co-expression of TNF and IFN-γ (**Fig. 2G, S2I**). Not only were the proportions of TNF^+^ T_CM_ cells higher following antigen stimulation of T cells from I.N. compared to S.C. vaccine groups, but higher expression of TNF was also apparent by flow cytometry (**Fig. 2H**). Stronger T_CM_ functional responses after I.N. vaccine administration is consistent with increased activity of memory cells that are able to home to multiple LNs by virtue of their CD62L expression^47^. These results support that, even controlling for vaccine formulation by using the same adjuvant, mucosal immunization promotes greater systemic antigen-specific memory responses compared to S.C. injection, where the response is concentrated in the draining LNs. Furthermore, recall of those T_MEM_ cells from I.N. vaccination leads to an enhanced polyfunctional phenotype based on the expression of multiple cytokines.

**Figure 2:**
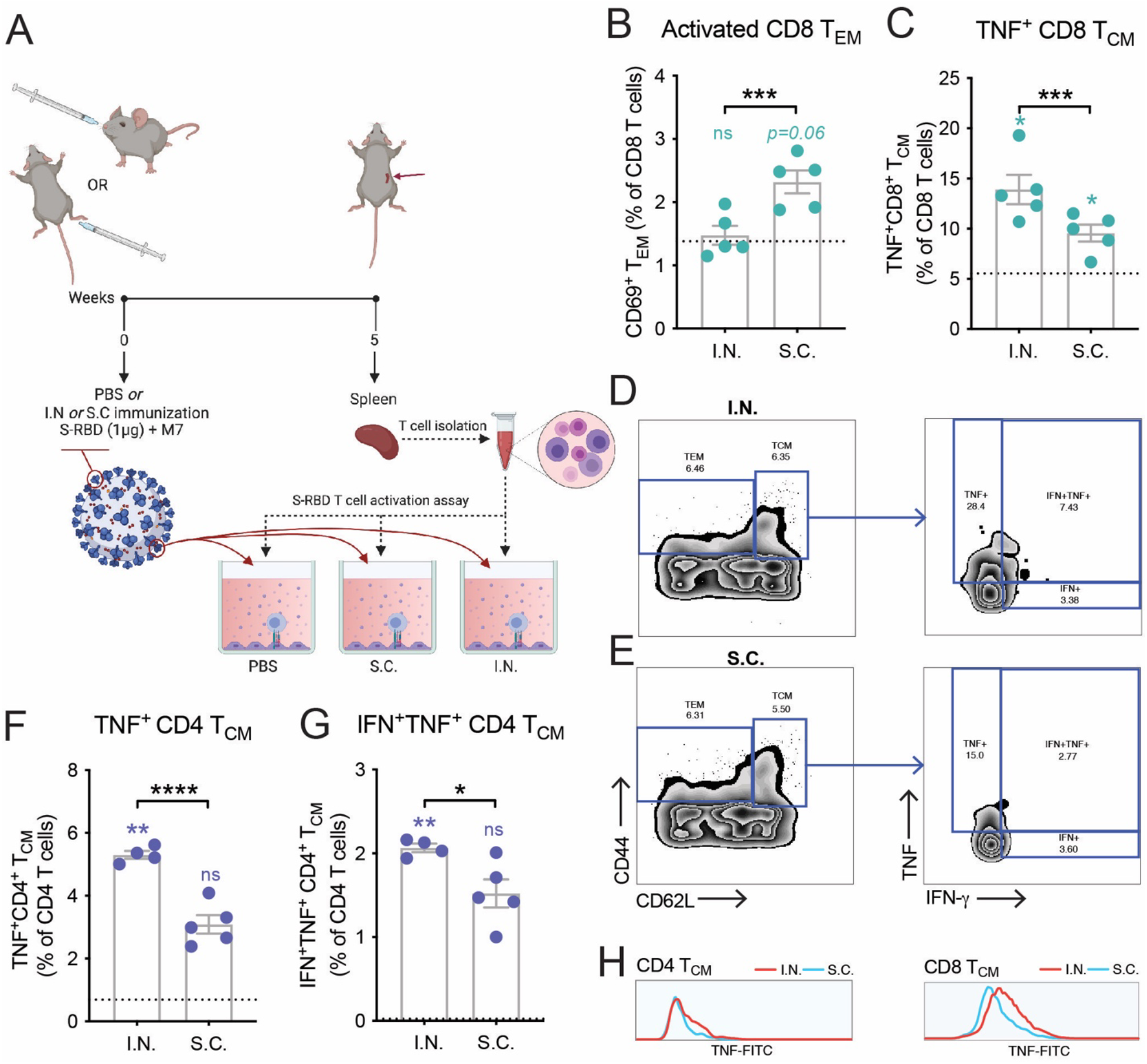
Mucosal vaccination enhances induction of antigen-specific polyfunctional T_CM_. (**A**) Schematic representing the experimental design where splenocytes isolated from mice vaccinated with M7+S-RBD by either the I.N. or S.C. route were stimulated with antigen S-RBD. (**B**) Increased activation of CD8 T_EM_ cells detected from mice vaccinated via the S.C. route, following stimulation with S-RBD. (**C**) Increased TNF^+^ CD8 T cells from mice vaccinated via the I.N. route following stimulation with S-RBD. (**D-E**) Representative flow cytometry plots showing the identification of T_EM_ and T_CM_ populations for the (**D**) I.N. and (**E**) S.C. vaccinated groups and their expression of cytokines IFN-γ and TNF, by intracellular staining. (**F-G**) Quantification of the (F) TNF^+^ and (G) IFN-γ^+^TNF^+^ populations of CD4 T_CM_ following antigen stimulation indicates an increase following I.N. compared to S.C. vaccination with the same M7+S-RBD. For, B-C and F-G, data points represent experimental replicates (individual donor mice) and dashed line represents the average baseline control for T cells from naïve mice stimulated with S-RBD. Significance compared to naïve controls is represented with colored symbols above each bar. (**H**) Representative histograms showing strong induction of TNF following I.N. compared to S.C. vaccination. *p<0.05, ***p<0.001 ****p<0.0001 by Student’s unpaired t-test.

### Improved polyfunctional memory recall by T cells upon challenge following mucosal vaccination

We next questioned whether the improved polyfunctional T cell phenotype induced by mucosal vaccination was T cell-intrinsic and if it would influence the character of immune activation upon antigen challenge. To address this question, we performed an adoptive transfer experiment using the Thy1.1/1.2 system for tracking donor versus recipient T cells (**Fig. 3A**). For this, T cells were harvested from the spleens of Thy1.2^+^ mice given I.N. or S.C. vaccination with M7+S-RBD, with a boost to ensure robust responses, and adoptively transferred into recipient mice with Thy 1.1^+^ T cells. To simulate a viral infection without the potential of differential viral replication kinetics influencing the T cell recruitment and activation, mice were given an I.N. challenge of full-length S protein. After allowing 5 days for memory recall and CD8 T cell trafficking, we isolated the lungs and their draining LNs, the brachial LNs, to assess the phenotypes and activation profiles of donor Thy1.2^+^ T cells (**Fig. 3A**, gating strategy in **S3A**). As expected, Thy1.2^+^ donor T cells could be detected in the lungs following S-antigen challenge (**Fig. 3B**) and these were mostly CD8 T cells (**Fig. 3C**), whereas CD4 cells in the lungs had very low frequency (**Fig. S3B**). Given the scarcity of donor CD4 T cells in lungs and based on the important functions of CD8 T cells in responding to viral infections in peripheral tissues, we focused on characterizing donor memory CD8 T cells. First, we noted there were no significant differences in the numbers of memory CD8 T cells that were recruited into the lung tissue between groups whose donor T cells were derived from I.N. versus S.C. vaccinated animals (**Fig. 3C**). However, we observed that donor memory CD8 T cells expressed higher levels of TNF compared to recipient memory CD8 T cells for both vaccination groups, with the highest levels expressed in the group having donor T cells from mice with I.N. vaccination (**Fig. 3D**), suggesting improved antigen-specific activation at the challenge site. The donor memory CD8 T cells in the lungs also expressed higher levels TNF as determined by MFI by flow cytometry (**Fig. 3E**) and, interestingly they also expressed higher levels of the activation marker CD69 by MFI (**Fig. 3F**). These results suggest that while there was no effect on the efficiency of memory CD8 T cell recruitment into the lungs following challenge, the T_MEM_ cells from I.N. vaccinated mice were more activated and functional following their entry into the lung tissue, supporting improved antigen-specific recall.

**Figure 3:**
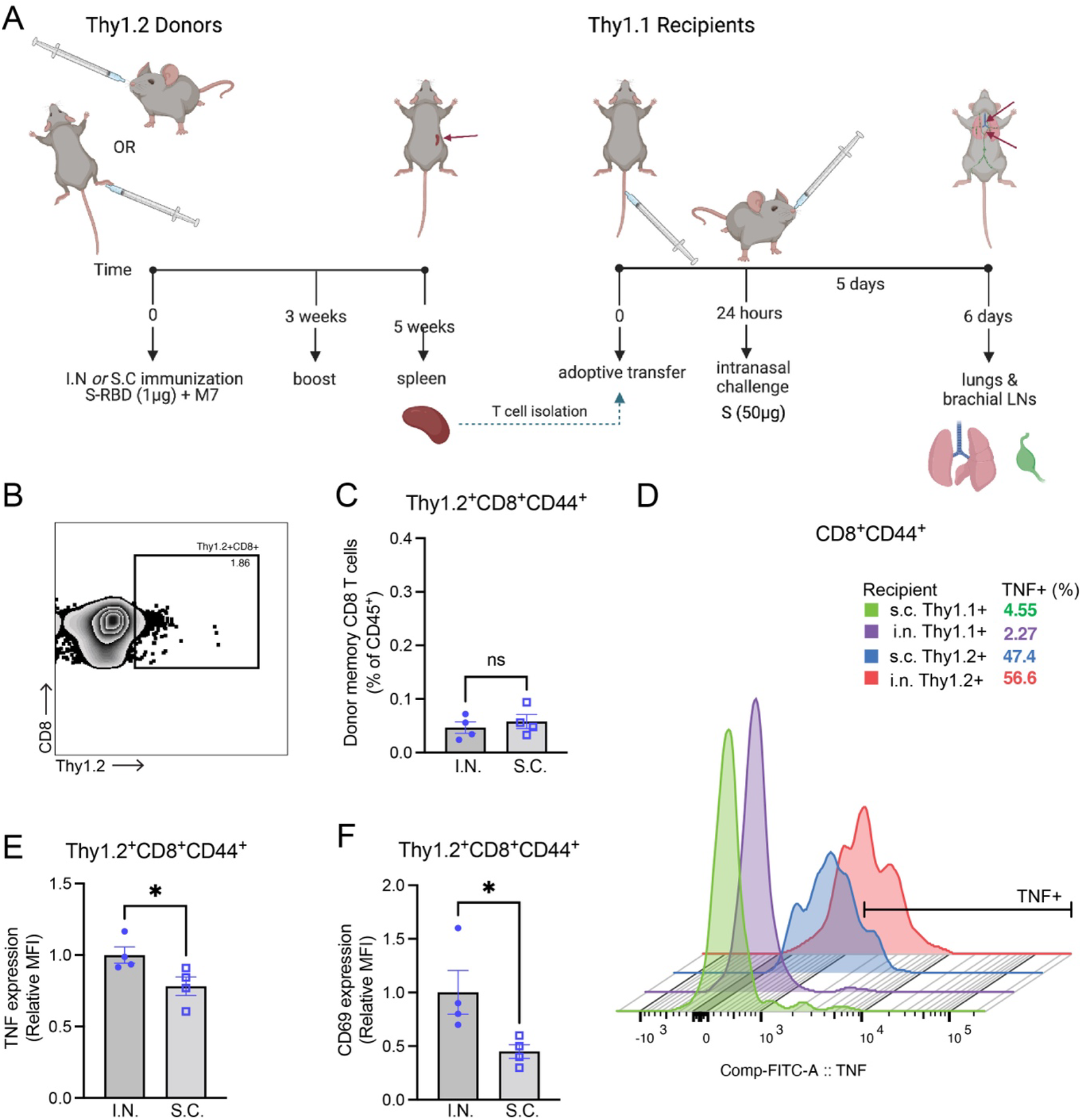
Improved re-activation of memory T cells derived from I.N. vaccinated donors in recipient lungs upon challenge. (**A**) Diagram illustrating the experimental design of adoptive transfer of purified Thy1.2^+^ T cells from M7+S-RBD-vaccinated donors into Thy1.1^+^ naïve recipients, followed by I.N. challenge with S protein. (**B**) Flow cytometry plot indicating the presence of donor Thy1.2^+^CD8^+^ T cells in the lungs of recipient mice 5 days after challenge. Plot depicts concatenated samples from all groups. Full gating strategy provided in **Fig. S3A** (**C**) Donor Thy1.2^+^CD8^+^ T cells constituted a minor portion of hemopoietic cells in the lung following challenge and did not differ in frequency between I.N. or S.C. vaccinated groups. (**D**) Histogram of TNF expression on donor (Thy1.2^+^) and recipient (Thy1.1^+^) CD44^+^CD8^+^ T cells representative of each group indicates an increase in TNF expression by donor T cells in lungs from vaccinated mice. Downsampling of 600 T cells was used to facilitate comparisons of equal numbers of cells for each sample. The percentage of TNF^+^ cells of total CD44^+^CD8^+^ T cells is indicated in the legend. (**E-F**) The MFI for (**E**) TNF expression and (**F**) CD69 expression were compared for Thy1.2^+^ donor CD8 T_MEM_ cells in the lungs following S challenge. *p<0.05 by Student’s unpaired t-test.

We also questioned whether S antigen challenge in animals (as performed in **Fig. 3A**) would induce heightened T_CM_ responses and polyfunctional T cell responses *in vivo* following transfer of T cells from I.N. compared to S.C. challenges, similar to our observations in *ex vivo* assays (**Fig. 2**). Donor T cells could be identified in the brachial LNs based on Thy1.2 expression (**Fig. 4A**). We examined the CD4 and CD8 subsets of memory donor T cells (**Fig. 4B**) and noted the presence of both CD4 and CD8 populations that expressed cytokines including TNF and IFN-γ (**Fig. 4C, S3D-E**) which were present in the LNs of mice that received donor T cells from I.N. but not S.C. vaccinated mice. Additionally, significantly increased proportions of donor CD8 T cells having the T_CM_ and T_EM_ phenotypes were present in brachial LNs for the I.N. vaccinated group (**Fig. 4D-E**). Furthermore, more CD8^+^ and CD4^+^ donor memory cells in the brachial LN produced cytokines TNF and IFN-γ following antigen challenge in mice receiving T cells derived from I.N. vaccination compared to S.C. vaccination (**Fig. 4F-G**). These results indicate that T cells develop an improved polyfunctional phenotype following I.N. vaccination, able to respond to challenge in the lung-draining LNs with higher levels of activation and cytokine production.

**Figure 4:**
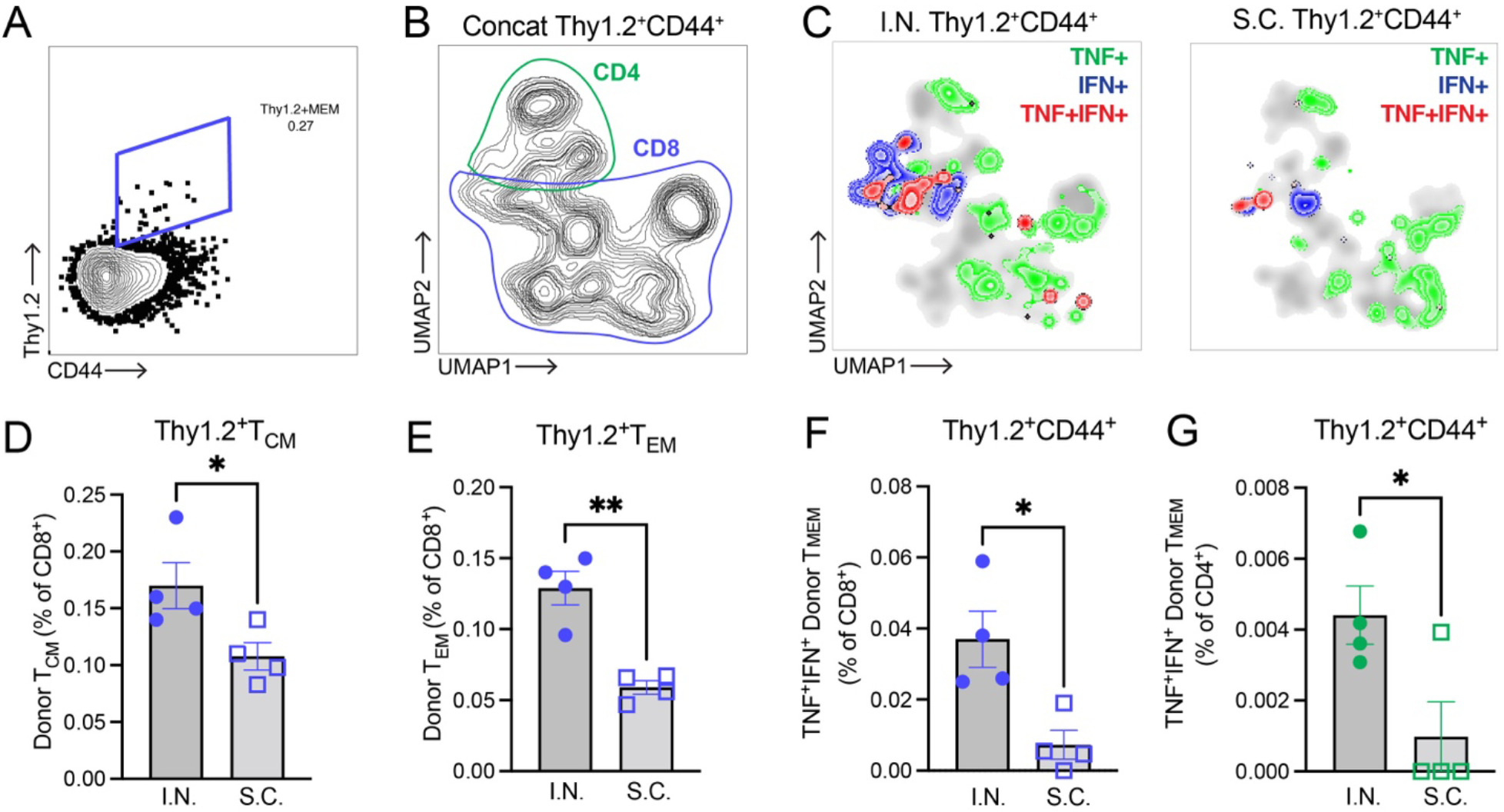
Mucosal vaccination enhances memory T cell responses in brachial LNs. (**A**) Donor Thy1.2^+^ T_MEM_ cells were detected in the brachial lymph nodes of recipient mice, as shown for a representative recipient. Full gating strategy is provided in **Fig. S3C**. (**B**) UMAP presentation of donor T_MEM_ cells (concatenated representation of all groups) with the locations of CD4 and CD8 T cell subsets outlined on the plot. (**C**) Donor T_MEM_ cells (Thy1.2^+^CD44^+^) expressing the cytokines TNF and/or IFN-γ are shown by overlaying the cytokine-positive subpopulations over the total UMAP plots for each group I.N. (left) vs. S.C. (right). (**D-E**) Plots indicating the percentage of donor-derived CD8 T cells with the (**D**) T_CM_ or (**E**) T_EM_ phenotypes. Percentage of (**F**) CD8 and (**G**) CD4 T cells that are donor T_MEM_ cells, staining double-positive for cytokines (TNF^+^IFN-γ^+^). *p<0.05, **p<0.01 by Student’s unpaired t-test.

### Superior antibody responses following mucosal vaccination

We identified improved systemic T cell responses following I.N. compared to peripheral S.C. vaccination; however, given the high correlation between antibody responses and protection against symptomatic infection for humans vaccinated against SARS-CoV-2^15^, we also questioned whether mucosal vaccination influenced antibody responses. To characterize these antibody responses, we first measured anti-S-RBD antibody titers by ELISA. Early responses at 3 weeks post-immunization showed no difference in binding to the same antigen, S-RBD, between mice exposed to the M7-S-RBD vaccine via the I.N. or S.C. routes; however, by 5 weeks post-immunization, anti-S-RBD titers were significantly higher in mice vaccinated via the mucosal route (**Fig. 5A**). No significant differences in antibody avidity were observed to the same S-RBD used as the vaccine antigen (**Fig. 5B**), suggesting the polyclonal antibodies had a similar polyclonal strength of binding to the original antigen used for vaccination. We also wanted to gain an understanding of whether the antibodies induced could be protective against SARS-CoV-2 and, therefore, used a surrogate virus neutralization test which detects total immunodominant neutralizing antibodies targeting S-RBD^48^. The surrogate neutralization test against S-RBD from an ancestral strain similar to the antigen used for vaccination, the Singapore/2/2020 strain, revealed that there was effective and similar concentration-dependent induction of neutralizing antibodies by both I.N. and S.C. vaccination routes (**Fig. 5C**). We also investigated the potential of antibodies generated by I.N. versus S.C. vaccine exposure to induce antibodies with cross-neutralizing capacity towards other variants of SARS-CoV-2. While S.C. inoculation induced efficient neutralizing antibodies against the Alpha variant in addition to the parental strain, cross-neutralization against other variants tested was low for the Delta, Delta plus, and gamma VOCs and not significantly present for others (**Fig. 5D, S4**). In contrast, I.N. vaccination induced more broadly cross-protective antibodies, with more significant neutralization at higher dilutions, which was significant compared to naïve controls for all variants tested, including Alpha, Delta, Beta, Gamma, Delta plus, Lambda, Mu, OmicronBA.1, and OmicronBA.2 (**Fig. 5D, S4**). This suggests that mucosal vaccination alters the breadth of neutralizing antibodies and may promote broadly neutralizing responses.

**Figure 5:**
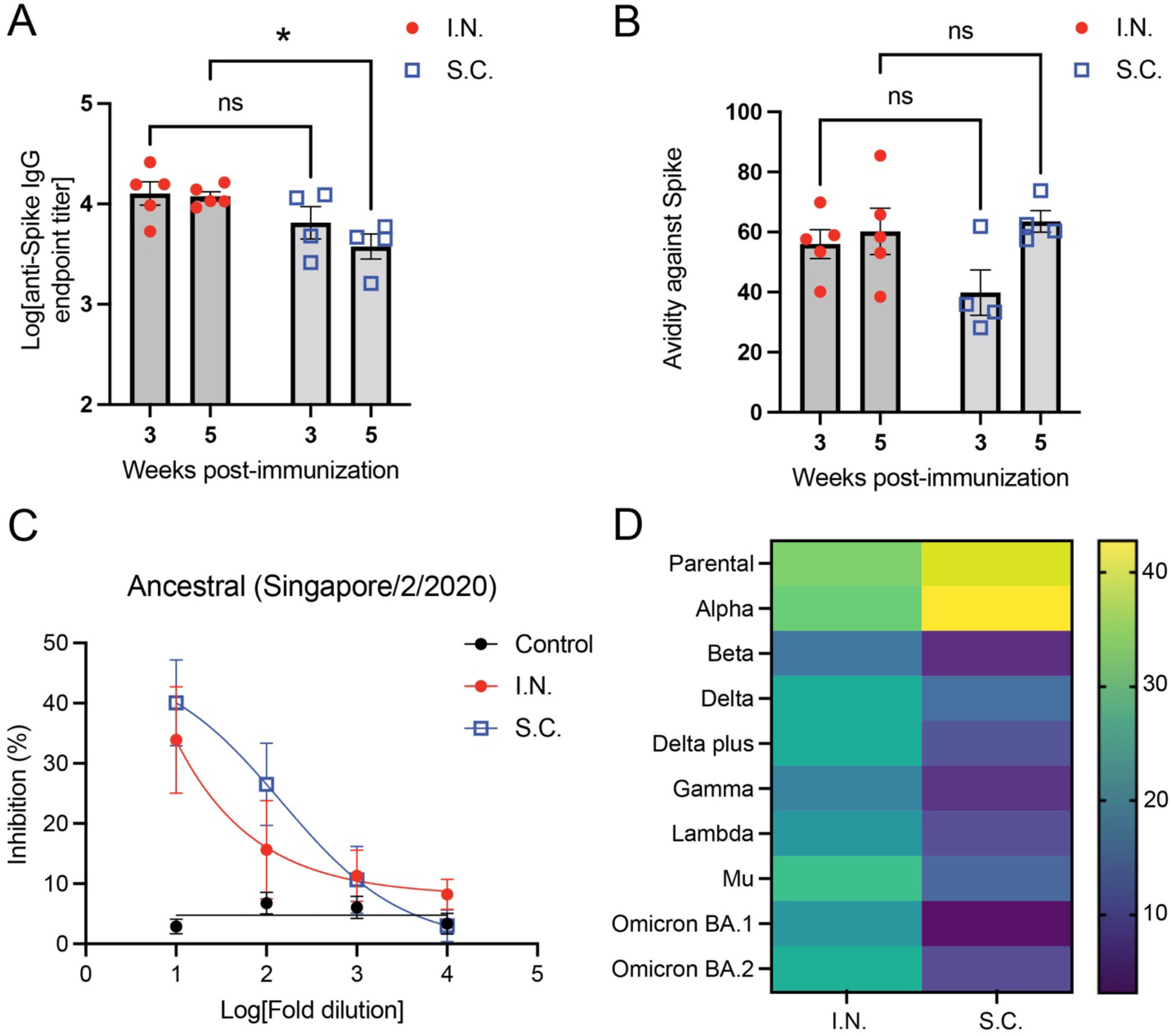
Superior antibody titer and SARS-CoV-2 variant cross-neutralization after mucosal vaccination. (**A**) Anti-S-RBD endpoint titers and (**B**) Avidity (percentage antibody that remains bound after stringent ELISA washing) following S.C. or I.N. vaccination. *p<0.05, by two-way ANOVA. ns=not significant. (**C**) Percentage inhibition of S-RBD association with its receptor hACE-2 by s-VNT. For control vs. I.N, p=0.003; for control versus S.C. p<0.001. For I.N. versus S.C., the comparison was not significantly different. (**D**) Heatmap depicting the % inhibition against S-RBD from multiple SARS-CoV-2 variants at 1:10 serum dilution. Corresponding dose response curves with p-values are provided in panel **C** and **Fig. S4**.

## Discussion

Mucosal vaccination has been increasingly acknowledged as a promising strategy to improve next-generation COVID-19 vaccine efficacy and immune protection of the airways^37,49-54^. Here we show that mucosal vaccination against S-RBD antigen of SARS-CoV-2 promotes a T cell-intrinsic phenotype that is associated with superior systemic immune responses and antibody responses that result in improved antibody persistence *in vivo* and cross-protection against SARS-CoV-2 variants compared to S.C. vaccination with the same formulation.

We observed the heighted systemic T cell responses in multiple T cell subsets in the spleen during the acute activation phase following mucosal vaccination, which also persisted in the T_MEM_ cell compartment several weeks following challenge. T_MEM_ cells generated by mucosal vaccination also exhibited an improved polyfunctional phenotype, characterized by dual expression of TNF and IFN-γ, both upon *ex vivo* stimulation as well as *in vivo* memory recall to antigen. *In vivo* splenic T cell responses during the acute phase following vaccination were also characterized by improved IL-17 production. Interestingly, IL-17 has been associated with lung IgA secretion^55^. Adoptive transfer of T cells obtained after resolution and contraction of the vaccine-induced response from vaccine recipients who had been given the exact same formulation of antigen and adjuvant, differing only by site of inoculation, confirmed the T cell-intrinsic imprinting of the site of inoculation on the T_MEM_ phenotype. Importantly, improved T cell activation and polyfunctionality in the lungs following antigen-challenge occurred in recipient mice who had been transferred T cells from mucosal rather than S.C. vaccination. This occurred even though similar numbers of T cells were recruited into the lung tissue for both recipient groups. Consistent with our study, others have also observed robust T cell activation following mucosal vaccination in animal models of subunit vaccination with S protein^41^. Our work illustrates that the polyfunctional nature of T_MEM_ cells, independent of their peripheral tissue homing abilities, could contribute to mucosal site protection and highlights the role of T cells in establishing systemic mucosal vaccine-induced memory since these responses can be adoptively transferred by T cells. A goal of vaccination is to induce the type of immune response that most closely approximates natural immune protection against infection, while eliminating risks of disease. For COVID-19, despite some controversy^56^, human vaccinees to parenteral SARS-CoV-2 do not appear to induce robust airway-resident antigen-specific T cells, unlike those who experienced natural infections^57,58^. This highlights the potential of next-generation COVID-19 vaccines to improve mucosal and systemic immune responses through modulation of T cells.

Of the T_MEM_ cells affected by our vaccination strategy, we identified here that T_CM_ cells are a central component of systemic mucosal vaccine-induced immunity. Activated and/or polyfunctional T_CM_ cells were observed in the spleen following vaccination, as well as in the T_MEM_ compartment that was effectively recalled by antigen re-stimulation. While antigen-specific T_CM_ were also present in subjects that received S.C. administered vaccine, and they could also be reactivated following adoptive transfer, as expected, both their numbers and the magnitude of their cytokine production responses were heightened in mucosal vaccinees. T_CM_ cells are particularly defined by their expression of CD62L, which allows them to roll on high endothelial venules and enter secondary lymphoid tissues^59^. It is likely that the increased numbers of CD62L-expressing cells within the T_MEM_ compartment following mucosal vaccination is key for the homing of these cells to the spleen that typifies increased systemic immunity. These results were observed in the context of antigen delivery with an appropriate mucosal adjuvant, while equivalent concentrations of antigen alone induced sub-optimal T cell activation responses. Efficient conversion to the T_CM_ phenotype could allow broader dissemination of antigen-specific T_MEM_ cells, which could increase the chances of subsequent exposure to antigen and memory recall and/or potential to provide B cell help in other lymphoid organs. During memory recall, T_CM_ are also thought to serve as a pool of T cells that can replenish the T_EM_ population^60-63^ and, consistent with this, *in vivo* antigen challenge was also associated with significantly increased numbers of CD8 T_EM_ in the lung-draining brachial LNs of recipients of T cells from I.N. vaccinated groups, even though *ex vivo* antigen restimulation of spleen T cells prior to transfer resulted in improved activation for CD8 T_EM_ in the S.C. vaccinated group. These results highlight the potential of site-specific immune responses to influence the long-term balance of T cell sub-populations.

Systemic T cell responses are also likely to impact B cell dependent antibody responses, owing to the influence of T cells, particularly CD4 T cells, on B cell help and germinal center activity^64,65^. A subset of CD4 T_CM_ expressing CXCR5 are also highly consequential to germinal center production since they can upregulate BCL-6 during memory recall, promote plasma cell differentiation and drive secondary germinal center formation and antibody production^66-69^. Consistent with this, we have identified improved antibody responses that coincide with the characterization of mucosal vaccine-induced responses as dominated by polyfunctional T_CM_. Both S.C. and I.N. vaccination strategies induced S-specific antibodies of similar avidity and neutralization towards the parental S protein, yet differences in antibody responses were also significant. I.N. vaccination induced a small but significantly higher level of antibodies at 5 weeks after the final vaccine boost and more broadly neutralizing antibodies against multiple VOC, compared to S.C. vaccination. Since broadly neutralizing responses were induced without altered avidity, this suggests that the mucosal vaccination strategy might have resulted in the preservation of antibodies against more diverse epitopes within the polyclonal pool.

Our results here are unique in that we are able to directly compare the responses induced by the same dose of antigen for two routes of immunization since M7 adjuvant is effective as an adjuvant both when injected in the skin and also when administered at mucosal surfaces. However, the limitations of S.C. vaccination were not the result of the adjuvant alone, as M7 has been shown to perform well as an adjuvant when injected S.C.^34,36^, and these differences were also consistently observed when compared to the human-approved adjuvant, Alum. Even so, it is also possible that some of the vaccine induced effects observed here are adjuvant specific, since the adjuvant’s mechanism of mast cell activation (likely with some other beneficial effects on other myeloid cells^70^) is unique compared to other strategies, and since mast cell phenotypes in these tissues are different^71^.

Improved systemic immune responses and improved variant cross-neutralizing antibodies generated by mucosal vaccination could be applied to next generation vaccines against SARS-CoV-2 and other respiratory pathogens. Indeed, the current situation where vaccines rely on high titer specific neutralizing antibodies with limited induction of mucosal responses have room for improvements. Strategies of vaccination against SARS-CoV-2 with improved capacity to limit vaccine breakthrough infections and to reduce transmission are still needed and this study and others support that mucosal vaccination is a promising strategy to meet these goals.

## Methods

### Animal studies

All animal studies were conducted at the vivarium in Duke-NUS Medical School and approved by the SingHealth IACUC. C57BL/6 mice purchased from InVivos were used for all experiments and immunizations began when they were 8-10 weeks old.

### Vaccinations

Mice were vaccinated with 1 μg of recombinant S-RBD protein (Sino Biological) either with or without 20μg of Mastoparan-7 (M7). For some groups, S-RBD was resuspended in alum. Footpads were injected with a 20μL volume of vaccine or vehicle control (PBS). For nasal inoculations, the same doses of 1 μg S-RBD + 20μg M7 were instilled in a volume of 12 μL per mouse (6 μL per nostril).

### Flow cytometry

NALT or popliteal LNs were harvested at necropsy along with spleens. The tissues were digested with collagenase (Sigma) and passed through 70μm cell strainers (Corning) to prepare single cell suspensions. RBCs were lysed to remove them from spleen single cells using RBC lysis solution (BioLegend). Total cell numbers were determined by counting on a hemocytometer. To facilitate intracellular staining for cytokines, single cell suspensions were incubated for 5 hours in 2μM monensin (BioLegend) to inhibit intracellular protein trafficking. Cells were stained with Live/Dead Fixable Blue Dead cell stain (Invitrogen, L23105) for 10 minutes prior to staining with anti–CD45-BUV395 (564279), anti–CD3e-PercP-Cy5.5 (551163), anti–CD4-BV650 (563232), anti–CD8a– Alexa Fluor 700 (557959), and anti– CD69-FITC (557392) (all from BD Biosciences), anti–NK1.1-PE (eBioscience, 12-5941-82), and anti–γδ TCR–APC (BioLegend, 118116) for 1h. Subsequently, 3x with 1% BSA in PBS solution, fixed with 4% paraformaldehyde (PFA) for 20 mnutes on ice, and permeabilized with 0.1% saponin in 1% BSA in PBS solution. Intracellular staining was done for IFN-γ (anti-IFN-γ-APC-Cy7, BioLegend, 505850) and IL-17a (anti-IL-17A-BV510, BD Biosciences, 564168) for 1 hr. Cells were washed 3x with 0.1% saponin in 1% BSA-PBS solution and finally resuspended in 1% BSA-PBS. Cells were acquired using a LSRFortessa cell analyzer (BD Biosciences) and analysed using FlowJo software (version 10). Heatmaps were generated using Heatmapper^72^ after normalization to saline-challenged controls and log-transformation of data.

### ELISA

Recombinant S-RBD protein (Sino Biological) was coated onto 96 well plates in carbonate (15mM) bicarbonate (35mM) buffer at 4°C, overnight. Serial 2x dilutions of serum were added to the coated plates and incubated overnight at 4°C. Plates were washed 3x with PBS and treated with an AP conjugated anti-mouse IgG antibody (Southern Biotech) for 1.5 hours. For avidity ELISA, plates were washed with 4M urea for 10 minutes prior to addition of secondary antibodies. Plates were washed again 3x with PBS and AttoPhos substrate (Promega) was added to each well. Fluorescence intensity at excitation/emission 440/560nm was measured using a Tecan Spark 10M plate reader after 45 minutes.

### Surrogate virus neutralization assay

A study team member blinded to the experimental groups performed the multiplex sVNT assay as previously described^48,73^. In brief, serum samples were pre-incubated with avidin microspheres coated with AviTag-biotinylated RBD proteins from different SARS-CoV-2 strains (including ancestral, Alpha, delta, Lambda, Beta, Gamma, Mu, Delta plus, Omicron BA.1 and BA.2) for 15 min at 37°C, followed by addition of Phycoerythrin (PE)-labelled human ACE2 at a final concentration of 2,000 ng/ml for another 15 min at 37°C. After two washes, the signals were acquired using the Luminex MAGPIX ireader.

### T cell activation assay

JAWSII cells (1.5×10^4^) were seeded to each well of a 96-well flat-bottom plate in αMEM with 5 ng/mL GM-CSF and incubated at 37°C with 5% CO_2_ in atmospheric air. 24 hours after seeding 1μg of S-RBD protein was added to each well. On day 3 post-seeding, spleens were harvested from vaccinated mice at day 35 post-vaccination and prepared as described above. Wells were washed with PBS and 1×10^5^ splenocytes were added to each well in RPMI containing 10% FBS. Cells were incubated at in a 5% CO_2_ incubator at 37°C for four days before analysis by flow cytometry. At day 7 post-seeding, cells were incubated for 5 hours in 2μM monensin (BioLegend) to inhibit intracellular protein trafficking and then stained with Live/Dead Near IR Dead cell stain (Invitrogen, L10119) for 10 minutes. Cells were then stained with anti–CD3e-PercP-Cy5.5 (551163), anti–CD4-BV650 (563232), anti–CD8a–Alexa Fluor 700 (557959), anti-CD44-BV510 (563114), anti-CD62L-PE-Cy7 (560516, all from BD Biosciences) and anti–CD69-eFluor450 (11-0691-82, eBioscience) for 1 hour in PBS supplemented with 1% BSA on ice. Subsequently, cells were washed 3x with 1%BSA in PBS, fixed with 4% PFA at 4°C for 20 minutes, and permeabilized with 0.1% Saponin in 1% BSA-PBS solution for 30 minutes. Intracellular staining was done for IFN-γ (anti-IFN-γ-BV711, BD Biosciences 554412) and TNFα (goat-anti-TNFα, R&D Systems AF-410-NA, anti-goat-IgG-FITC, Jackson ImmunoResearch, 305-096-006) for 1 hour in permeabilization solution. Data were acquired using LSRFortessa cell analyzer (BD Biosciences) and analysed using FlowJo software (version 10).

### T cell adoptive transfer and antigen challenge

Spleens were harvested from vaccinated mice at day 35 post vaccination and single cell suspensions were prepared as described above. RBCs were lysed using 1x RBC lysis solution (BioLegend) and cells were counted using a hemocytometer. T cells were isolated from the splenocytes using Pan T Cell Isolation Kit II (Miltenyi BioTec) according to the manufacturers protocol and 1×10^6^ T cells were transferred to Thy1.1 mice by tail vein injection. Mice were challenged with 50μg S protein (full length) intranasally in 20μl PBS 24h post-transfer. The SARS-CoV2 S protein was expressed using the vector pCAGGS containing the SARS-Related Coronavirus 2, Wuhan-Hu-1 Spike Glycoprotein Gene from BEI (NR-52394) and purified following the published protocol^74^. Mice were euthanized 5 days post-challenge and lungs and brachial lymph nodes were harvested. The tissues were digested with collagenase (Sigma) and passed through 70μM cell strainer (Corning) to prepare single cell suspensions. RBC lysis was done for spleen single cells using RBC lysis solution (BioLegend). Cells were stained with Live/Dead Blue Dead cell stain (Invitrogen, L23105) for 10 minutes. Cells were then stained with anti–CD45-BUV395 (564279), anti–CD4-BV650 (563232), anti–CD8a–Alexa Fluor 700 (557959), and anti-CD44-BV510 (563114), anti-CD62L-PE-Cy7 (560516, all from BD Biosciences), anti–NK1.1-PE (eBioscience, 12-5941-82), anti–CD69-eFluor450 (eBioscience, 11-0691-82, or PerCP-Cy5.5 45-0691-82) anti-CD90.2-PerCP-Cy5.5 (BioLegend, 140322, or eFluor450 eBioscience 48-0902-80) and anti–γδ TCR–APC (BioLegend, 118116). Subsequently, cells were washed, fixed with 4% paraformaldehyde (PFA), and permeabilized with 0.1% Saponin in 1% BSA-PBS solution. Intracellular staining was done for IFN-γ (anti-IFN-γ-BV711, BD Biosciences 554412) and TNFα (goat-anti-TNFα, R&D Systems AF-410-NA, anti-goat-IgG-FITC, Jackson ImmunoResearch 305-096-006) in permeabilization solution for 1 hour on ice. Cells were acquired with a LSRFortessa cell analyzer (BD Biosciences) and analysed using FlowJo software (version 10).

### Statistical analysis and data presentation

Data were analysed using Prism 9 and Microsoft Excel software. For multiple groups comparison, ANOVA was used, while Student’s unpaired t-test was used when two groups were compared. All data are presented as the means of experimental replicates using individual mice and error bars represent the SEM throughout the manuscript. All data obtained are included in the manuscript with the exception of one flow cytometry sample that was determined to have poor cell viability by live-dead staining. Figures were prepared using Adobe Illustrator and diagrams were drawn using biorender.com.

## Acknowledgements

The authors thank Antonio Bertoletti for insightful discussions. This study was funded by Duke-NUS start-up funding and Singapore Ministry of Education funding to ALS (MOE-T2EP30120-0011) and National Medical Research Council funding to LFW (STPRG-FY19-001, COVID19RF-003, COVID19RF-060 and OFLCG19May-0034).

## Notes

### Competing Interest Statement

The authors have declared no competing interest.

