## Supplemental Figures for "Mucosal vaccination for SARS-CoV-2 elicits superior systemic T central memory function and cross-neutralizing antibodies against variants of concern"

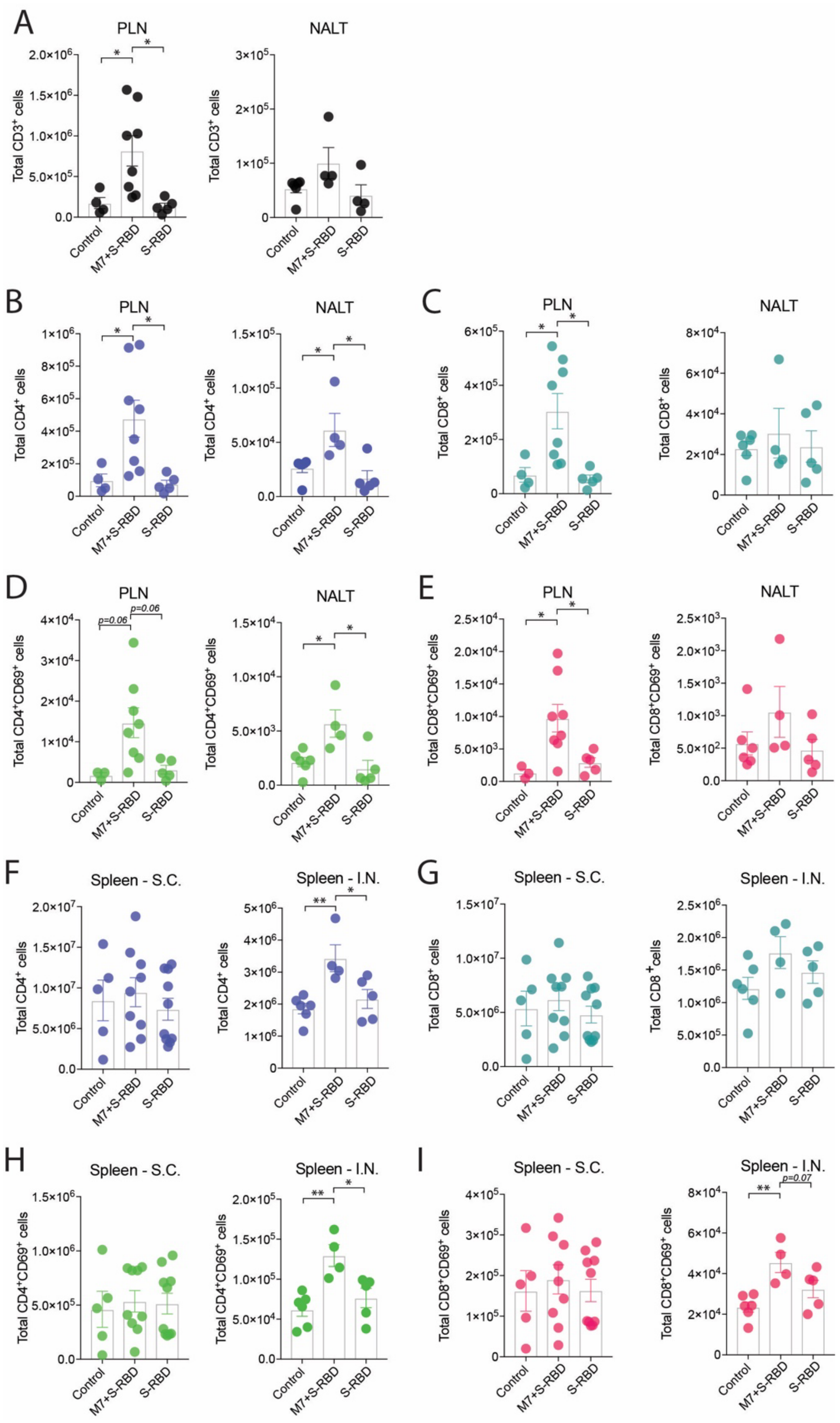


**Figure S1: Superior systemic T cell activation following mucosal vaccination**
Plots showing the total (**A**) T cells, and (**B**) $CD4^+$ (**C**) $CD8^+$ (**D**) activated $CD4^+$ or (**E**) activated $CD8^+$ T cells in the draining lymphoid tissue of the vaccine administration, either PLN or NALT, day 5. Plots showing the total (**F**) $CD4^+$ (**G**) $CD8^+$ (**H**) activated $CD4^+$ (**I**) activated $CD8^+$ T cells in the spleen day 5 following vaccination. Groups were compared by 1-way ANOVA with Holm-Sidak's posttest; $^*p<0.05$; $^{**}p<0.01$. Non-significant p-values less than 0.1 are indicated on the graph.

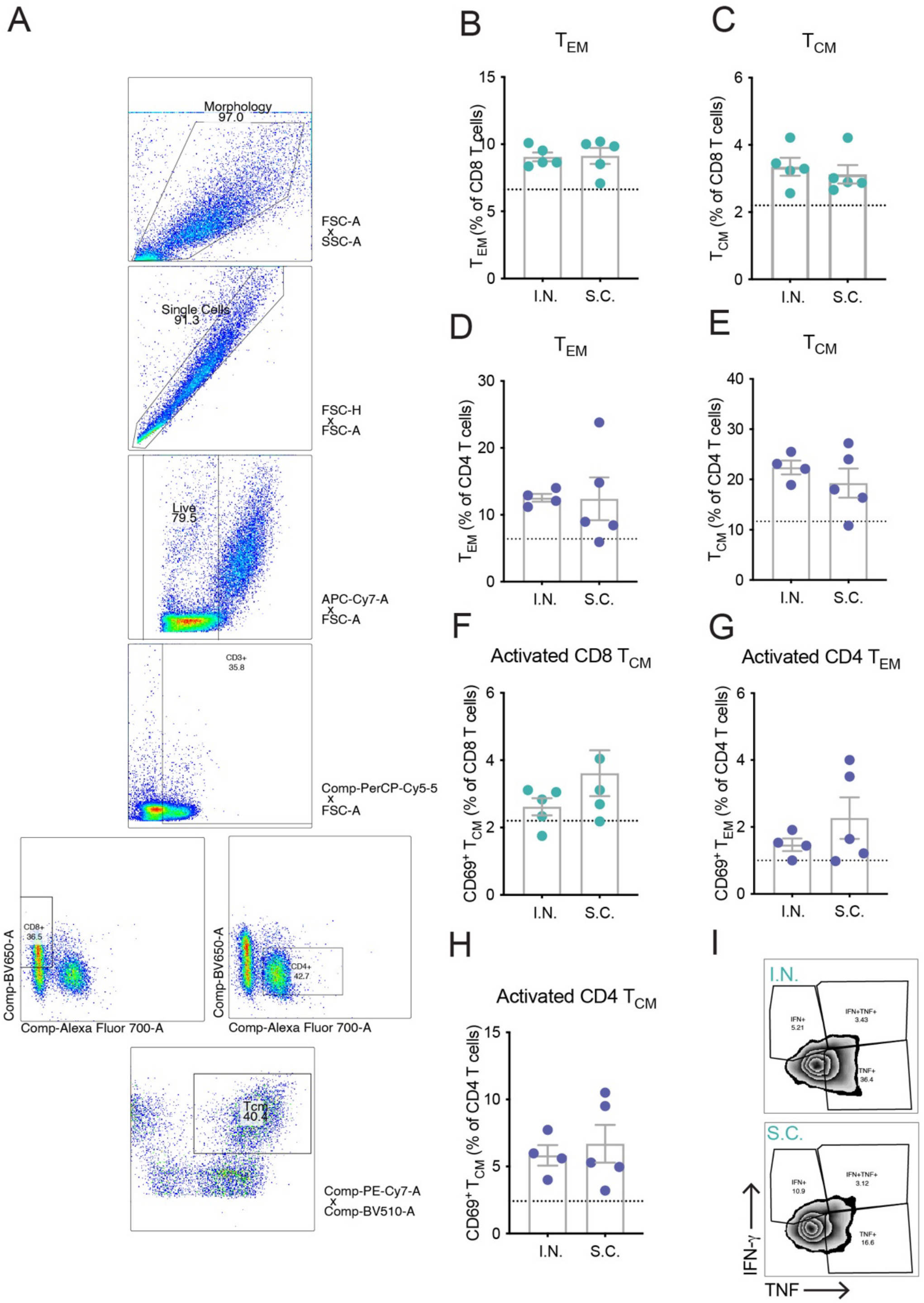


**Figure S2: Effective activation of several $T_{MEM}$ subsets by *ex vivo* antigen stimulation**
(**A**) Gating strategy to identify memory T cell subsets in co-culture. Percentages of (**B-C**) CD8 memory and (**D-E**) CD4 memory subsets or (**F**) Activated CD8 $T_{CM}$ or activated (**G-H**) memory CD4 T cells, following S-RBD stimulation. All groups in B-H were compared by Student's unpaired t-test and found to not differ significantly. (**I**) Representative plots showing intracellular staining of $T_{MEM}$ cells for IFN-γ and TNF for I.N. and S.C. groups.

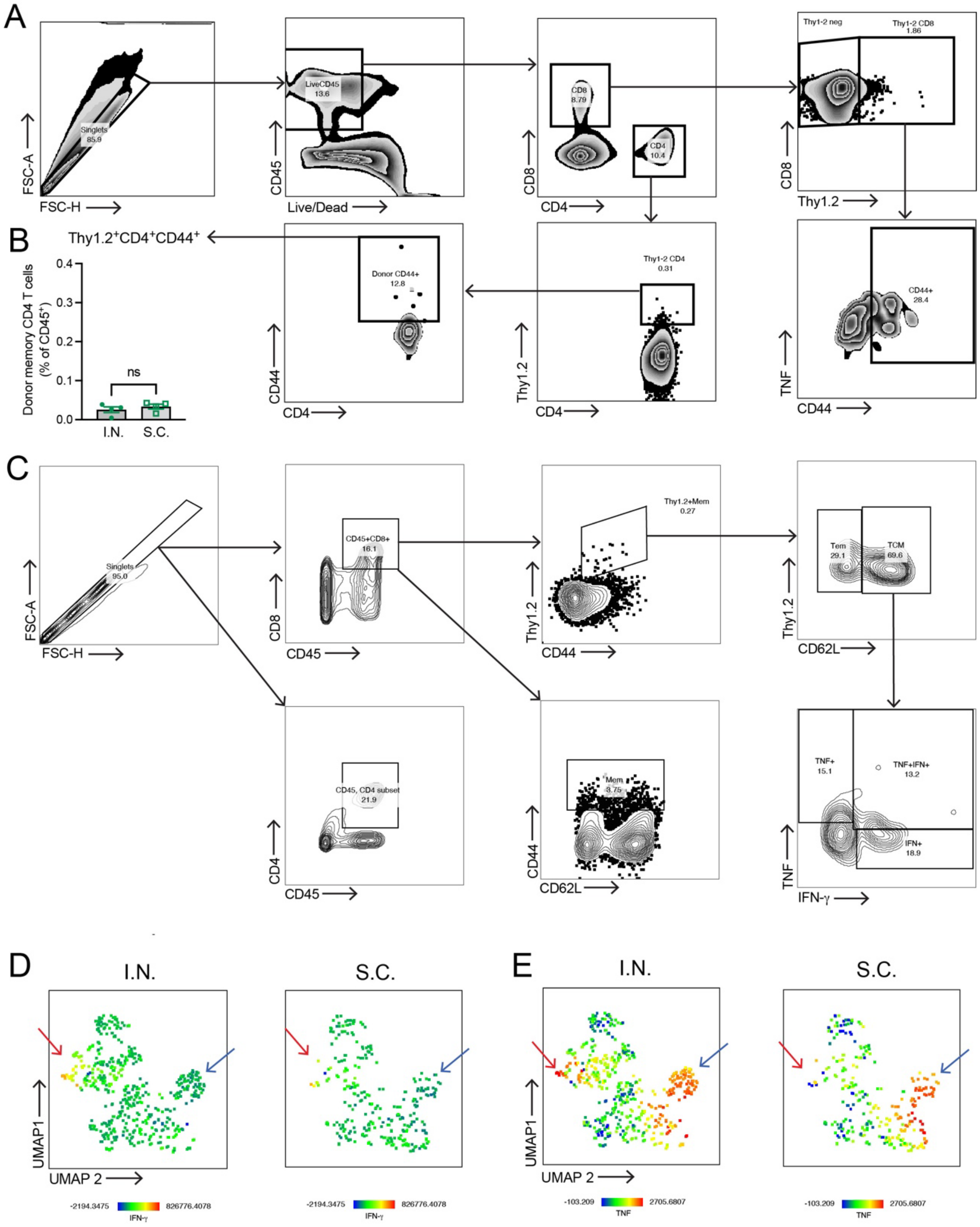


**Figure S3. Identification and characterization of donor T cell phenotypes following *in vivo* antigen challenge**

(**A**) Flow cytometry gating strategies to identify donor Thy1.2$^+$ T cell populations in the lungs (**B**) Donor Thy1.2$^+$ CD4 T cells constituted a minor portion of hemopoietic cells in the lung following challenge and did not differ in frequency between I.N. or S.C. M7+S-RBD vaccinated groups. (**C**) Flow cytometry gating strategies to identify donor Thy1.2$^+$ T cell populations in the brachial LNs. (**D-E**) Heat maps depicting (**D**) IFN-γ and (**E**) TNF expression in $T_{MEM}$ cells of the brachial LNs. Areas with cytokine expressing CD4 T cells are indicated with red arrows and areas with cytokine expressing CD8 T cells are indicated by blue arrows.

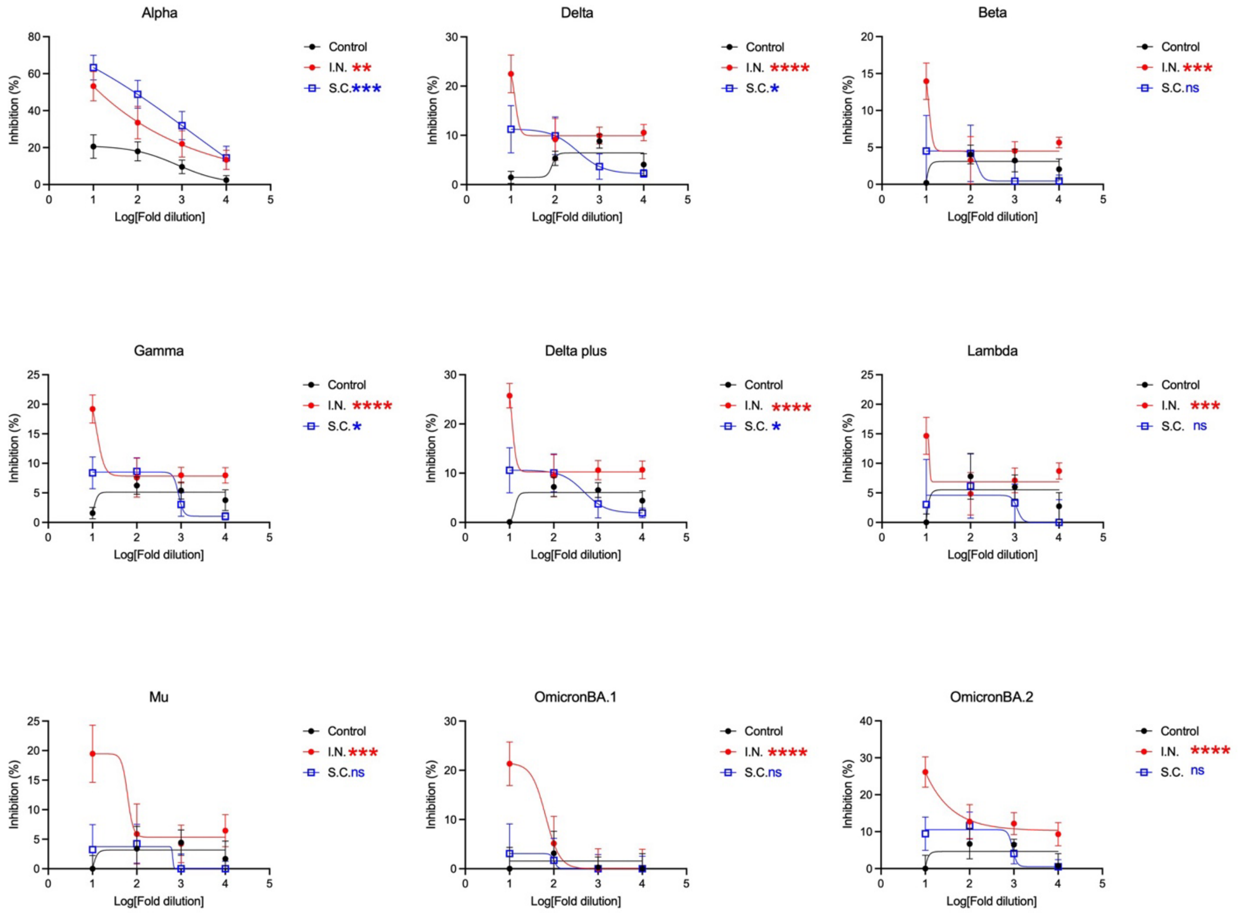


**Figure S4. Superior serum cross-neutralization of SARS-CoV-2 variants following mucosal immunization.** Dose response curves showing the % inhibition against S-RBD from multiple SARS-CoV-2 variants by s-VNT assay. P-values were determined by 2-way ANOVA. *p<0.05, **p<0.01, ***p<0.001 ****p<0.0001 by two-way ANOVA.

**Table S1: Predicted mouse MHC-I-binding peptides in the RBD of S-protein**

Peptide predictions for binding to the MHC-I molecules found in C57Bl/6 mice (H-2-Db, H-2-Dd, H-2-Kb and H-2-Kd) for the RBD domain of S-protein (Accession # QHD43416) were performed using TepiTool[1] and the top 2% of predicted binders based on percentile rank are presented.

| Peptide start | Peptide end | Peptide | IEDB consensus percentile rank | Allele |
|---|---|---|---|---|
| 126 | 134 | KVGGNYNYL | 0.09 | H-2-Db |
| 48 | 56 | SVLYNSASF | 0.1 | H-2-Db |
| 187 | 195 | YQPYRVVVL | 0.17 | H-2-Db |
| 47 | 56 | YSVLYNSASF | 0.17 | H-2-Db |
| 7 | 17 | SIVRFPNITNL | 0.17 | H-2-Db |
| 9 | 17 | VRFPNITNL | 0.25 | H-2-Db |
| 159 | 168 | STPCNGVEGF | 0.3 | H-2-Db |
| 64 | 72 | VSPTKLNDL | 0.33 | H-2-Db |
| 125 | 134 | SKVGGNYNYL | 0.37 | H-2-Db |
| 8 | 17 | IVRFPNITNL | 0.38 | H-2-Db |
| 179 | 187 | FQPTNGVGY | 0.58 | H-2-Db |
| 191 | 199 | RVVVLSFEL | 0.65 | H-2-Db |
| 192 | 200 | VVVLSFELL | 0.68 | H-2-Db |
| 90 | 100 | RQIAPGQTGKI | 0.69 | H-2-Db |
| 207 | 215 | CGPKKSTNL | 0.72 | H-2-Db |
| 115 | 123 | VIAWNSNNL | 0.78 | H-2-Db |
| 127 | 134 | VGGNYNYL | 1.1 | H-2-Db |
| 64 | 72 | VSPTKLNDL | 0.01 | H-2-Dd |
| 207 | 215 | CGPKKSTNL | 0.01 | H-2-Dd |
| 206 | 215 | VCGPKKSTNL | 0.02 | H-2-Dd |
| 187 | 195 | YQPYRVVVL | 0.04 | H-2-Dd |
| 205 | 215 | TVCGPKKSTNL | 0.05 | H-2-Dd |
| 185 | 195 | VGYQPYRVVVL | 0.07 | H-2-Dd |
| 92 | 100 | IAPGQTGKI | 0.13 | H-2-Dd |
| 208 | 215 | GPKKSTNL | 0.2 | H-2-Dd |
| 185 | 193 | VGYQPYRVV | 0.2 | H-2-Dd |
| 51 | 59 | YNSASFSTF | 0.25 | H-2-Dd |
| 9 | 17 | VRFPNITNL | 0.29 | H-2-Dd |
| 64 | 74 | VSPTKLNDLCF | 0.3 | H-2-Dd |
| 2 | 11 | VQPTESIVRF | 0.43 | H-2-Dd |
| 10 | 17 | RFPNITNL | 0.46 | H-2-Dd |
| 6 | 14 | ESIVRFPNI | 0.49 | H-2-Dd |
| 185 | 192 | VGYQPYRV | 0.51 | H-2-Dd |
| 62 | 72 | YGVSPTKLNDL | 0.58 | H-2-Dd |

| | | | | |
|---|---|---|---|---|
| 129 | 137 | GNYNYLYRL | 0.01 | H-2-Kb |
| 7 | 14 | SIVRFPNI | 0.01 | H-2-Kb |
| 193 | 200 | VVLSFELL | 0.01 | H-2-Kb |
| 131 | 138 | YNYLYRLF | 0.03 | H-2-Kb |
| 185 | 192 | VGYQPYRV | 0.04 | H-2-Kb |
| 185 | 193 | VGYQPYRVV | 0.06 | H-2-Kb |
| 9 | 17 | VRFPNITNL | 0.07 | H-2-Kb |
| 116 | 123 | IAWNSNNL | 0.1 | H-2-Kb |
| 192 | 200 | VVVLSFELL | 0.14 | H-2-Kb |
| 24 | 32 | FNATRFASV | 0.15 | H-2-Kb |
| 49 | 56 | VLYNSASF | 0.2 | H-2-Kb |
| 192 | 199 | VVVLSFEL | 0.2 | H-2-Kb |
| 64 | 72 | VSPTKLNDL | 0.2 | H-2-Kb |
| 127 | 137 | VGGNYNYLYRL | 0.2 | H-2-Kb |
| 130 | 137 | NYNYLYRL | 0.21 | H-2-Kb |
| 136 | 143 | RLFRKSNL | 0.21 | H-2-Kb |
| 127 | 134 | VGGNYNYL | 0.25 | H-2-Kb |
| 186 | 194 | GYQPYRVVV | 0.05 | H-2-Kd |
| 61 | 69 | CYGVSPTKL | 0.08 | H-2-Kd |
| 32 | 40 | VYAWNRKRI | 0.11 | H-2-Kd |
| 176 | 185 | SYGFQPTNGV | 0.19 | H-2-Kd |
| 9 | 17 | VRFPNITNL | 0.26 | H-2-Kd |
| 77 | 84 | VYADSFVI | 0.26 | H-2-Kd |
| 186 | 195 | GYQPYRVVVL | 0.27 | H-2-Kd |
| 10 | 17 | RFPNITNL | 0.3 | H-2-Kd |
| 60 | 69 | KCYGVSPTKL | 0.44 | H-2-Kd |
| 59 | 69 | FKCYGVSPTKL | 0.52 | H-2-Kd |
| 187 | 195 | YQPYRVVVL | 0.56 | H-2-Kd |
| 81 | 89 | SFVIRGDEV | 0.61 | H-2-Kd |
| 50 | 59 | LYNSASFSTF | 0.68 | H-2-Kd |
| 186 | 193 | GYQPYRVV | 0.7 | H-2-Kd |
| 185 | 194 | VGYQPYRVVV | 0.74 | H-2-Kd |
| 154 | 162 | IYQAGSTPC | 0.78 | H-2-Kd |
| 31 | 40 | SVYAWNRKRI | 0.86 | H-2-Kd |

**Table S2: Predicted mouse MHC-II-binding peptides in the RBD of S-protein**

Peptide predictions for binding to the MHC-II molecule found in C57Bl/6 mice (H2-IAb) were performed using TepiTool[1].

| Peptide start | Peptide end | Peptide sequence | Consensus percentile rank | Allele |
|---|---|---|---|---|
| 194 | 208 | VLSFELLHAPATVCG | 2.9 | H2-IAb |
| 151 | 165 | STEIYQAGSTPCNGV | 3.7 | H2-IAb |
| 45 | 59 | ADYSVLYNSASFSTF | 5.5 | H2-IAb |
| 56 | 70 | FSTFKCYGVSPTKLN | 6.45 | H2-IAb |
| 27 | 41 | TRFASVYAWNRKRIS | 7.45 | H2-IAb |
| 173 | 187 | PLQSYGFQPTNGVGY | 7.55 | H2-IAb |

1 Paul, S., Sidney, J., Sette, A. & Peters, B. TepiTool: A Pipeline for Computational Prediction of T Cell Epitope Candidates. *Curr Protoc Immunol* **114**, 18 19 11-18 19 24, doi:10.1002/cpim.12 (2016).
